# Telemetry reveals rapid duel-driven song plasticity in a naturalistic social environment

**DOI:** 10.1101/803411

**Authors:** Pepe Alcami, Shouwen Ma, Manfred Gahr

## Abstract

Singing by songbirds is a complex motor skill learnt during juvenile development or, in ‘open-ended’ learners, before the onset of the breeding season. Outside of these specific periods, it is believed that birdsong does not change. Challenging this, here we demonstrate that in a seasonal songbird, social interactions during the breeding season induce a novel form of singing plasticity in naturalistic social environments. Using custom-made telemetric backpack technology to monitor song-based communication from freely-behaving canaries, we show that males temporally overlap their songs during aggressive ‘duels’. Singing duels induce an unexpected fast plasticity in song length, thereby enhancing singing performance and flexibility of a sexually-selected behavior. Remarkably, dueling canaries sing acoustically-similar songs, suggesting that competition within a specific song acoustic space drives dueling behavior. Overall, our findings reveal a previously unrecognized type of song plasticity different from the well-studied slow song plasticity as an imitation process for display purposes.

## Introduction

A key component of natural environments in which animals and their brains have evolved is the social context in which individual animal behavior takes place. Social environments have been shown to influence the production and fine-tuning of learnt motor skills (S. L. King, Sayigh, Wells, Fellner, & Janik, 2013; Lefebvre & Giraldeau, 1996; Ota, Gahr, & Soma, 2018; Svensson & Sheldon, 1998). The interaction of animals through vocal communication, a complex motor skill, embeds them in social networks that are critical to understand individual vocal behavior (McGregor, Butlin, Guilford, & Krebs, 1993). Singing by songbirds is one such motor skill influenced by con-specifics (Tchernichovski, Lints, Mitra, & Nottebohm, 1999). In songbird species where only males sing, females listening to singing males as well as songs from other males influence singing behavior (Beecher, Campbell, Burt, Hill, & Nordby, 2000; A. P. W. King, Meredith J., 1989; Payne, 1983; Sossinka & Böhner, 1980). In fact, singing by songbirds is involved in a plethora of social functions, including mate attraction, pair bond formation, parental investment, territorial behavior and signaling of social hierarchy (Akçay, Tom, Campbell, & Beecher, 2013; Collins, 2004; Ficken, Ficken, & Witkin, 1978; Garcia-Fernandez, Amy, Lacroix, Malacarne, & Leboucher, 2010; Kroodsma, 1976; McGregor et al., 1993). However, whether songs change during natural counter-singing interactions of songbirds in social environments remains widely unexplored.

Songs from songbirds are fine tuned throughout learning until they become stable or ‘crystallized’. In ‘close-ended’ songbirds like zebra finches (*Taeniopygia guttata*), songs crystallize during development. In contrast in seasonal songbirds like canaries (*Serinus canaria*), songs become plastic yearly before they crystallize during each breeding season. As a result, songs strongly differ across seasons (Leitner, Voigt, Garcia-Segura, Hof, & Gahr, 2001; Nottebohm, Nottebohm, & Crane, 1986). Once song is crystallized, singing occurs with increased stereotypy in terms of temporal and spectral properties, syllable sequence structure and length (Alliende, Giret, Pidoux, Del Negro, & Leblois, 2017; Hahnloser, Kozhevnikov, & Fee, 2002). Canary songs therefore constitute a suitable model to study the impact of social contexts on the plasticity of ‘crystallized’ song.

The ecological, behavioral and evolutionary frameworks of singing interactions have been described in a number of species, where the timing and acoustic properties of songs from interacting birds relative to each other can encode aggressive information (Araya-Salas, Wojczulanis-Jakubas, Phillips, Mennill, & Wright, 2017; Dabelsteen, McGregor, Holland, Tobias, & Pedersen, 1997; Mennill & Ratcliffe, 2004; Naguib & Kipper, 2006; Vehrencamp, 2001). Most studies have however used song playbacks to study counter-singing, that is, songs played on a loudspeaker, and not in contexts in which birds vocally interact. They therefore lack the essential interactive component between singing birds. The study of directly-interacting birds has been hurdled by the necessity to disambiguate signals from individual birds, particularly when birds overlap their songs. A first attempt to overcome this difficulty consists in the use of microphone arrays (Araya-Salas et al., 2017). However, they lack the temporal and acoustic precision to characterize in greater detail singing interactions due to their distance to sound sources. An alternative solution to record individual vocalizations that lacks these caveats is the use of individual microphones mounted on each animal (Gill et al., 2016). This technique, that we opted for here, allows the precise monitoring of individual songs.

In canaries, song playbacks have revealed that female canaries prefer overlapping to overlapped playbacks (Leboucher & Pallot, 2004). Overlapping another canary’s song may be interpreted as a signal of dominance (Amy et al., 2008; Leboucher & Pallot, 2004). Furthermore, female canaries allocate greater yolk amounts in their eggs in response to overlapping songs, highlighting the importance of overlapping behavior for biological fitness (Garcia-Fernandez et al., 2010). Because of all these reasons, canaries are a model species for which the behavioural and ecological implications of singing overlaps are best studied. However, whether counter-singing and overlapping behavior occur in social groups of canaries, what are the relative dynamics of singing and whether they impact songs remains elusive. Here, we combine recordings in the field and in the laboratory. In the laboratory, we use microphones mounted on individual canaries. These allow us to record their songs with telemetry with high temporal precision (Gill et al., 2016) in mixed aviaries (with both males and females) and characterize for the first time the relative singing dynamics of male canaries and their impact on song properties.

## Results

### Multiple-microphone recordings in the field reveal temporally-overlapping songs

We first asked whether canaries overlap their songs in their natural environment. For that purpose, we recorded wild canaries during the breeding season in Pico island (Azores, Portugal). Recordings were simultaneously performed with two directional microphones oriented towards different directions in fields where canaries had been located. Song spectrograms revealed canary song temporal overlaps, as shown by overlapping songs from different canaries preferentially recorded in one or the other microphone (Fig. 1).

**Figure 1.**
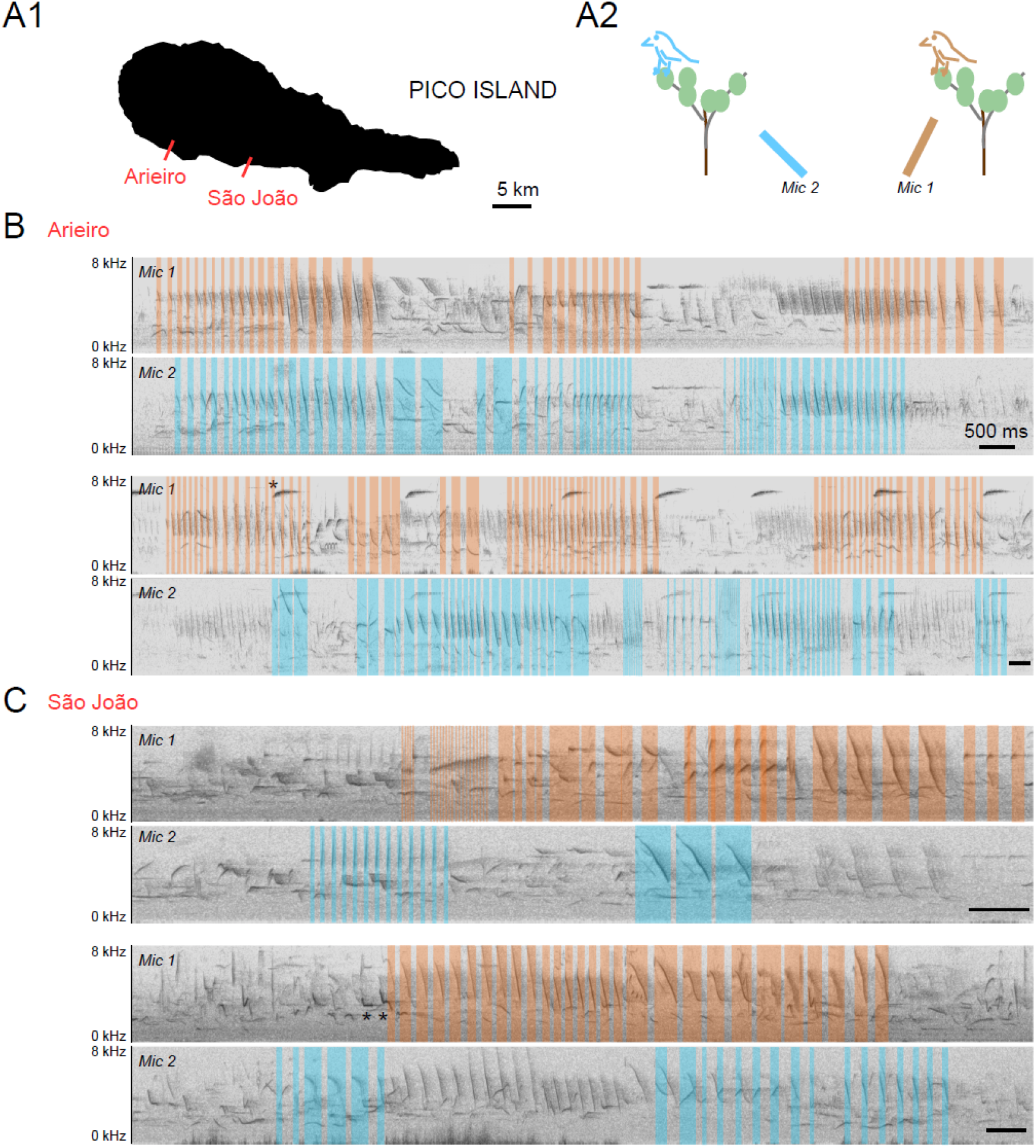
Multiple microphone recordings reveal overlapping songs from wild canaries. **A1.** Recording sites are indicated on a map of Pico island. **A2.** Recording configuration. Two directional microphones were directed to different directions in the field. **B.** Spectrograms of two example overlapping singing periods in Arieiro and **C.**, two example overlapping singing periods in São João. Syllables from putative canary songs from two different canaries recorded preferentially in microphone 1 (Mic1) or microphone 2 (Mic 2) are labelled in red and blue respectively. Scale bars, 500 ms. Note that recordings also show additional distant canary songs (top), and the vocal production of additional species (some examples being annotated with stars).

In the same fields where canaries could be recorded, the vocal production from a number of other bird species was also recorded. Frequently recorded species included chaffinches (*Fringilla coelebs ssp. moreletti*), blackbirds (*Turdus merula ssp. azorensis*), green finches (*Carduelis chloris*), blackcaps (*Sylvia atricapilla*) and robins (*Erithacus rubecula*). The vocal production of other bird species also happened to overlap canary songs, some of which have been labeled in Fig. 1.

In order to further characterize the intraspecific singing dynamics of male canaries with improved temporal detail and in better-controlled social environments that exclude inter-specific interactions, we designed a reduced social setting in aviaries. We kept canaries in breeding conditions with mixed sex groups formed by three males and two females, reproducing sex ratios previously reported in the wild (Voigt & Leitner, 1998). That is, we reproduced a competition context in which males compete for females during the breeding season.

### Telemetric recordings in naturalistic social environments reveal singing duels

We mounted custom-made ‘backpack microphones’ (Gill et al., 2016) on the back of freely-moving male domesticated canaries. This allowed us to record the individual vocal production simultaneously from all males with the help of a telemetric recording system (Fig. 2A,B).

**Figure 2.**
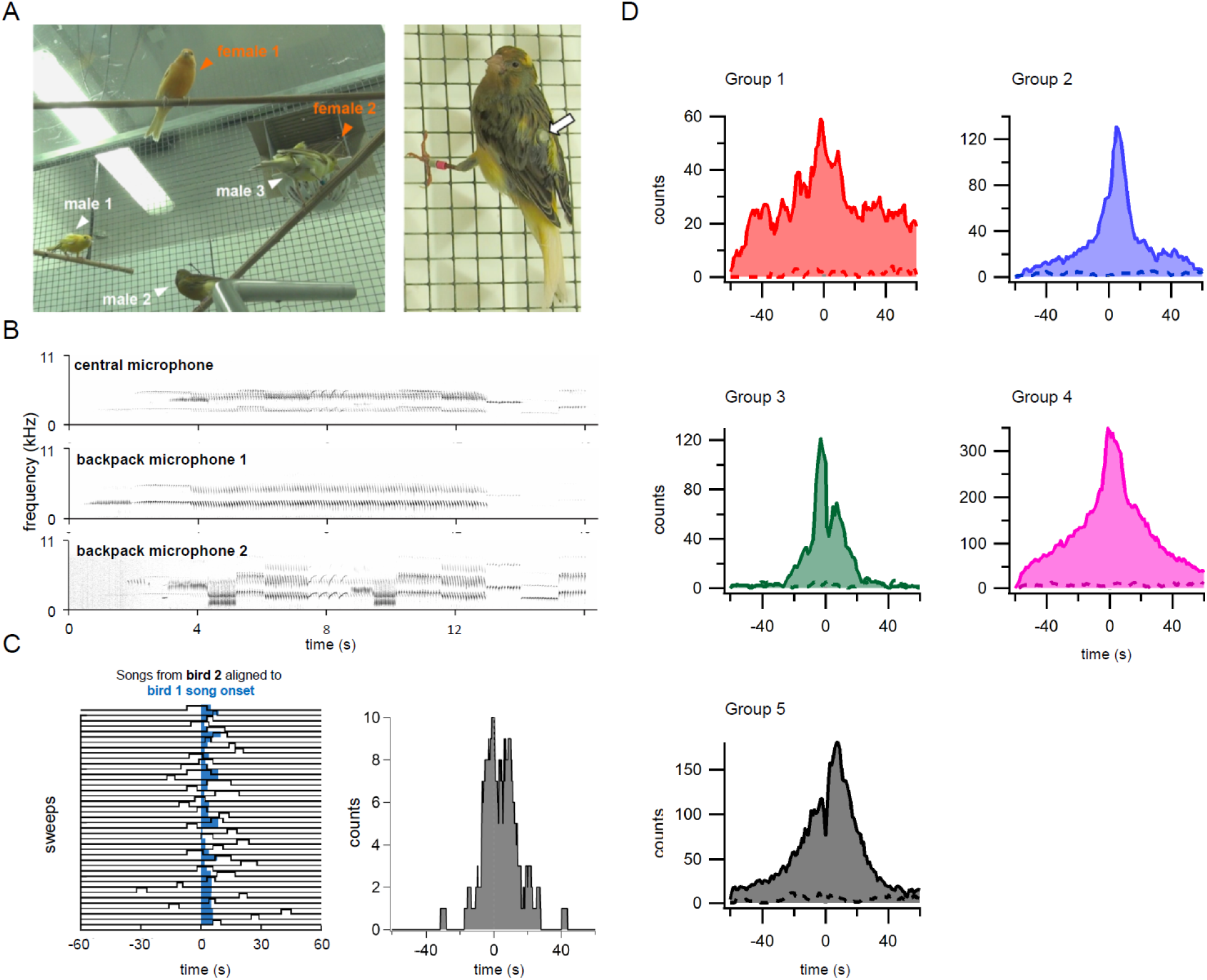
Telemetric recordings with backpack microphones reveal temporally-overlapping songs. **A.** Left, experimental design: mixed sex groups comprising two female and three male canaries, all three individually equipped with a backpack microphone. Right, male canary equipped with an individual microphone, pointed by the arrow. **B.** Representative simultaneous recording from a central microphone and from two backpack microphones confirms the identification of the individual vocal production by ‘backpack’ microphones. **C.** Left, example raster plot for all songs sung in a day by a canary (blue) and aligned vocal activity of a second canary (black). Right, corresponding histogram of the times at which songs were recorded in canary 2 centered at song onset from the reference canary 1. Bin size: 1 s. **D.** Data for all days in each group show increased singing when the singing activity of one bird was aligned to a second bird. Histograms were also generated from shuffled data (dashed lines), confirming that distributions show increased singing of canaries in the 60 s time-window relative to random overlaps. Bin size: 1 s.

Our results show that male canaries influence each other’s singing and tend to overlap their songs when they spontaneously interact with each other. Fig. 2C (left) illustrates the songs from a canary aligned to the song onset for all songs of another canary in the same aviary during one day. Fig. 2C (right) shows the corresponding histogram of the times at which songs were recorded centered at song onset from the reference canary. Inspection of the histogram reveals a bimodal distribution. The distribution at negative values in Fig. 2C corresponds to songs in which singing from the examined canary started before the reference canary started singing, whereas the distribution at positive values corresponds to songs occurring after the other canary started singing. In all five groups, and in 2 to 3 birds per group, the histogram of the times at which songs were recorded from one bird centered at song onset from another bird showed an increased singing activity relative to the histogram generated by shuffling randomly the onset of songs during the recording period (dashed lines in Fig. 2D).

Finally, in order to confirm the aggressive signaling nature of canaries singing together, we quantified the relationship between singing and physical fights from video recordings. Singing temporal overlaps were followed by fights in 51 ± 11 % of the cases, whereas individual singing preceded fights in only 13 ± 6 % of the cases (average from video recordings in n = 3 groups). All groups showed enhanced fighting associated to temporally-overlapping relative to solo singing (one-tail Fisher exact test; P < 0.05 in the three groups). Thus, song overlapping interactions are aggressive signals predictive of physical fights. We will henceforth call ‘duels’ the interactions in which canaries overlap their songs, influencing each other’s singing and establishing pairwise singing contests.

To summarize, a social competition context induces song overlapping between male canaries, influencing both the singing activity and the relative timing of singing. Moreover, duels are predictive of aggressive interactions.

### Plastic leader and follower dynamics

We observed frequent overlapping behavior from two male canaries in all groups, and occasionally from a third canary. The latter took more rarely part to duels, typically adopting a predominant soloist strategy, and potentially eventually joining and overlapping his songs with the group (Figs 3 and 4). Songs were recorded continuously for days, revealing overlapping behaviors in which both singers could lead or follow on a song-to-song basis. Remarkably, singing dynamics were plastic on a temporal scale of days, as can be appreciated in the three examples depicted in Fig. 3A1-A3.

**Figure 3.**
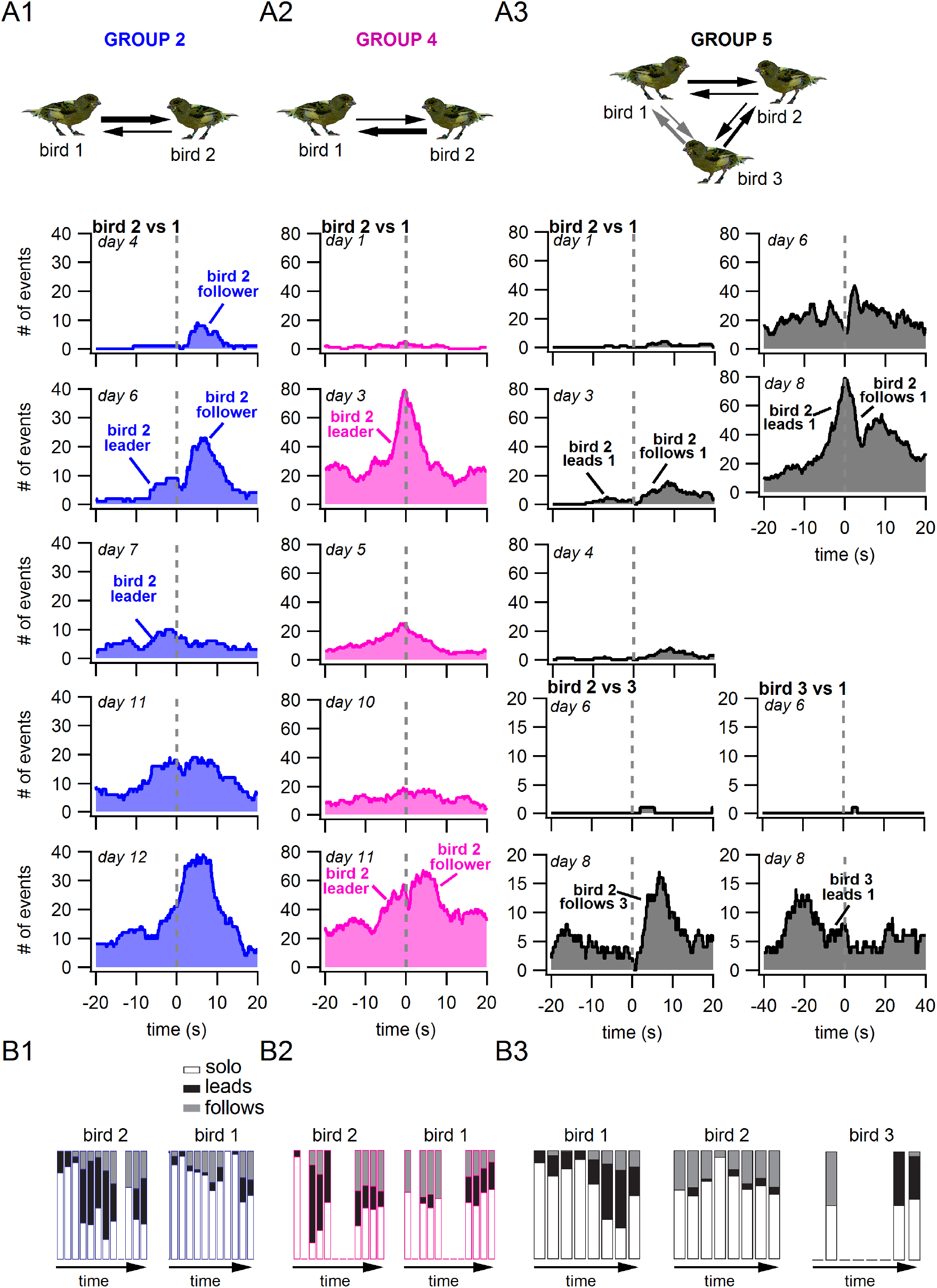
Plastic singing dynamics reveal temporally-evolving follower and leader roles. **A1, A2, A3.** Examples of singing interactions in three groups and their time course across several days. Note the change of scale in the bottom right histogram of the times at which songs were recorded centered at song onset from the reference bird. Bin size: 100 ms. **B1, B2, B3** illustrate the fraction of solo, leading and following songs in each group. Empty entries shown in figure 3B correspond to the absence of singing activity detected during the whole day. The color code for groups is the same as in Fig. 2.

**Figure 4.**
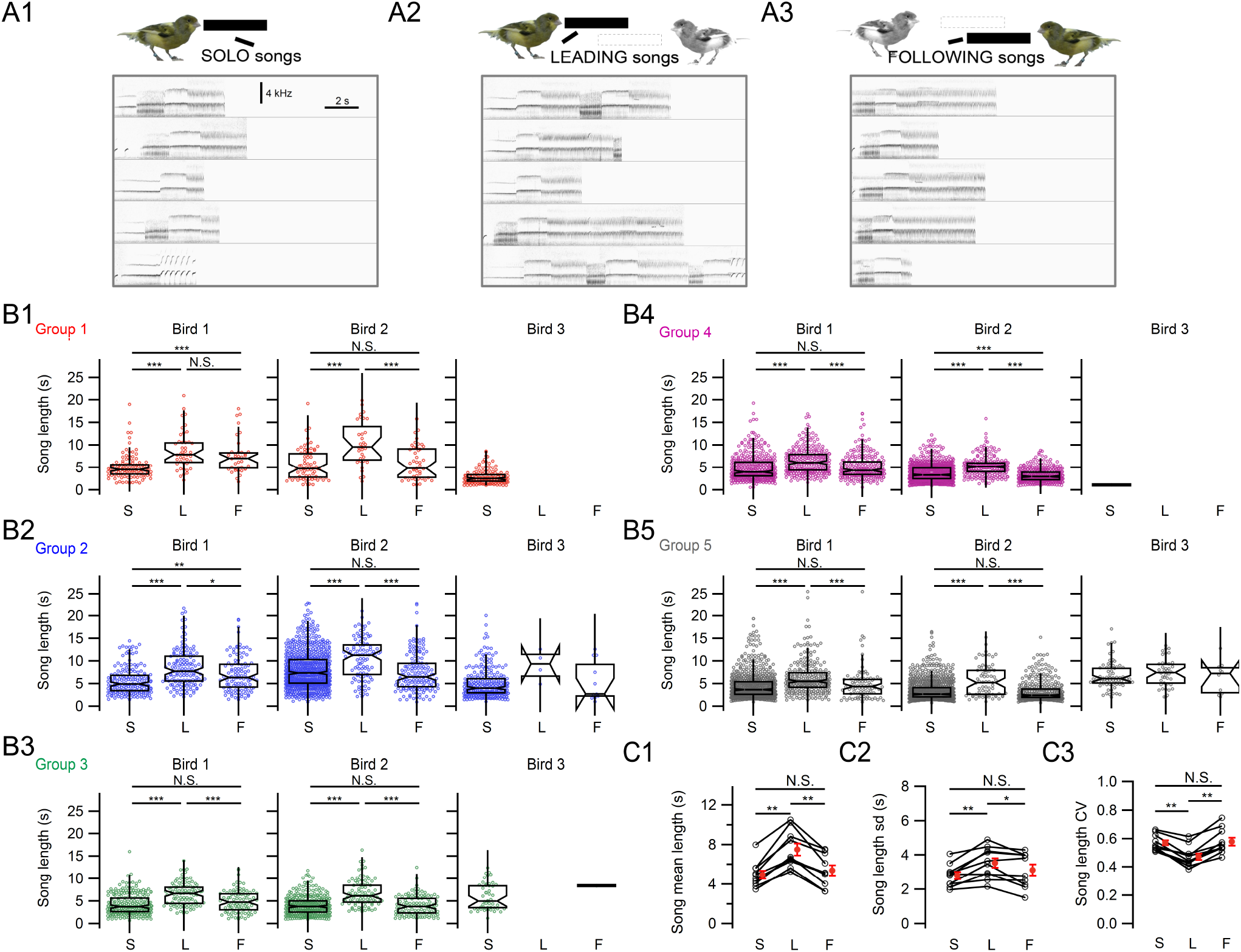
Song length and its variability depend on the relative timing of singing onset from male canaries. **A1-A3.** Representative examples of solo songs (A1), leading songs (A2) and following songs (A3). **B1-B5.**Song length for the three male canaries in each of the five groups were plotted in panels B1 to B5 corresponding to groups 1 to 5 shown in Fig. 2. The three different birds in a group are shown in the left, middle and right panel. S, solo songs; L, leading songs; F, following songs. Multi-comparison Dunn-Holland-Wolfe test for the two main duelists in each group, * P < 0.05, ** P < 0.01, *** P < 0.001. The color code for groups is the same as in Figs. 2 and 3. **C1.** Summary data show significant differences in average song length for solo, leading and following songs for ten birds in five groups. Song lengths from the same birds in different singing conditions are linked (S, solo songs; L, leading songs; F, following songs). * P < 0.05, ** P < 0.01, two-tail Wilcoxon Signed-Rank test. **C2.** Summary data show significant differences in the standard deviation of song length for solo, leader and follower songs. Open symbols, average song length from each individual bird. Filled symbols, average song length across all birds. * P < 0.05, ** P < 0.01, two-tail Wilcoxon Signed-Rank test. **C3.** Summary data show significant differences in the coefficient of variation of song length (CV) for solo, leader and follower songs. Open symbols, average song length from each individual bird. Filled symbols, average song length across all birds. * P < 0.05, ** P < 0.01, two-tail Wilcoxon Signed-Rank test.

The histogram of the times at which songs were recorded centered at song onset from the reference canary are shown in Fig. 3 for several groups during several days. The two birds from group 2 (Fig. 3A1, B1) start in a configuration in which bird 1 leads the singing duel and bird 2 follows. On the following days, leader/follower roles reconfigure: whereas bird 2 behaves as a follower during the first days, singing dynamics evolve to mixed leader/follower dynamics in which the two canaries both lead and follow. In group 4 (Fig. 3A2, B2) during the first days, singing activity of bird 2 shows a single mode of singing activity at short latencies hundreds of milliseconds relative to the reference bird 1 singing onset, mostly corresponding to a leader behavior: most songs from canary 2 start at negative time values before song onset of bird 1 and stop at positive values, after song onset of bird 1. With time, the distribution becomes wider. The temporal dynamics of group 5 (Fig. 3A3, B3) show plasticity in the timing of singing activity from bird 2 relative to bird 1: singing from bird 2 shifts from loosely following bird 1 (peak singing at 7.6 s-delay from song onset in bird 4) on day 1 to a reduced close-to-null latency between the peak of singing activity of bird 2 relative to the song onset of bird 1 on day 8. Interestingly, shortening of this latency occurred when the third male in the group, bird 3, also started to vocally interact with the two birds that initially overlapped their songs. Remarkably, bird 3 behaves as a leader, reconfiguring the relative singing dynamics in the whole group.

Overall, the relative timing of overlapping singing interactions in a group is variable and dynamic and evolves on a time scale of days.

### Leading, following and solo songs differ in length

We next examined the influence of the social singing context on songs in a song-to-song basis. We quantified song length in different overlapping conditions. Songs produced by the two canaries mediating most singing interactions in each group had a ~28 % longer duration during overlapping interactions than when the same birds sang solo songs (6.36 ± 0.55 s. vs 4.96 ± 0.41 s., P = 0.0019, paired Wilcoxon Rank test, n = 4084 solo songs, n = 2773 temporally-overlapping songs from n = 10 canaries from five groups).

Interestingly, song length differed depending on the relative timing of singing interactions: leading songs were on average longer than solo songs, systematically for all birds. In contrast, following songs were of different duration than solo songs in only three out of ten birds, being longer in two birds (Fig. 4A, B1-B5). These results reveal a common strategy to sing longer-lasting songs when canaries lead singing interactions, compared to a strategy only adopted by some of them to also sing longer songs when they behave as followers. Overall, across birds, leading songs were ~ 52 % longer than solo songs (7.53 ± 0.59 s vs. 4.96 ± 0.41 s, P = 0.0019, two-tail Wilcoxon Signed-Rank test, Bonferroni-corrected: P = 0.0059, n = 10 canaries, Fig. 4C1) and ~ 39 % longer than following songs (5.43 ± 0.59 s, P = 0.0019, two-tail Wilcoxon Signed-Rank test, Bonferroni-corrected: P = 0.0059, n = 10 canaries, Fig. 4C1). Solo songs were not significantly different to following songs, comparing mean lengths for all birds (P = 0.19, two-tail Wilcoxon Signed-Rank test, Bonferroni-corrected: P = 0.58, n = 10 canaries, Fig. 4C1).

We further examined whether the variability of song length was changing during singing interactions (Fig. 4C2). Remarkably, the standard deviation of song length also significantly increased by ~ 27 % for leading songs relative to solo songs (3.6 ± 0.3 s vs. 2.8 ± 0.2 s respectively, P = 0.0039, two-tail Wilcoxon Signed-Rank test, Bonferroni-corrected: P = 0.012, n = 10 canaries) and by ~ 13 % relative to following songs (3.1 ± 0.3 s, P = 0.014, two-tail Wilcoxon Signed-Rank test, Bonferroni-corrected: P = 0.041, n = 10 canaries). The standard deviation of solo and following song length was not significantly different (P = 0.11, two-tail Wilcoxon Signed-Rank test, Bonferroni-corrected: P = 0.32, n = 10 canaries). Thus, canaries sing songs that are more variable in length when they lead singing duels. We also examined the coefficient of variation as an additional measure of variability, normalized to the mean value (Fig. 4C3). Interestingly, the coefficient of variation of song length instead decreased by ~ 17 % and ~ 18 % for leading songs relative to solo and follower songs respectively (0.47 ± 0.02 vs. 0.57 ± 0.02 and 0.58 ± 0.03 respectively, P = 0.0020, two-tail Wilcoxon Signed-Rank test, Bonferroni-corrected: P = 0.0059, n = 10 canaries), and did not differ between solo and follower songs (P = 0.70, two-tail Wilcoxon Signed-Rank test, n = 10 canaries). Thus, despite song length variability increasing when birds lead duels, this variability decreases when scaled to the average song length.

In summary, song length and its variability depend on the relative timing in the singing interactions within a duel.

### Dueling canaries sing similar songs

We investigated the link between song properties and singing overlapping behavior by asking whether the songs sung by canaries that tend to duel in each group are more similar to each other than to the songs of the third canary, that we call ‘soloist’ since he rarely takes part to duels.

For this purpose, we analyzed songs sung in solo conditions from each canary to quantify the acoustic similarity between the songs from birds that tend to duel frequently relative to the third bird that rarely duels (Fig. 5). A detailed analysis of song features from the four groups for which we had enough songs to carry this analysis showed that syllable repetition rate, pitch and song length were more similar between the two birds that establish most duels in each group than with the soloist.

**Figure 5.**
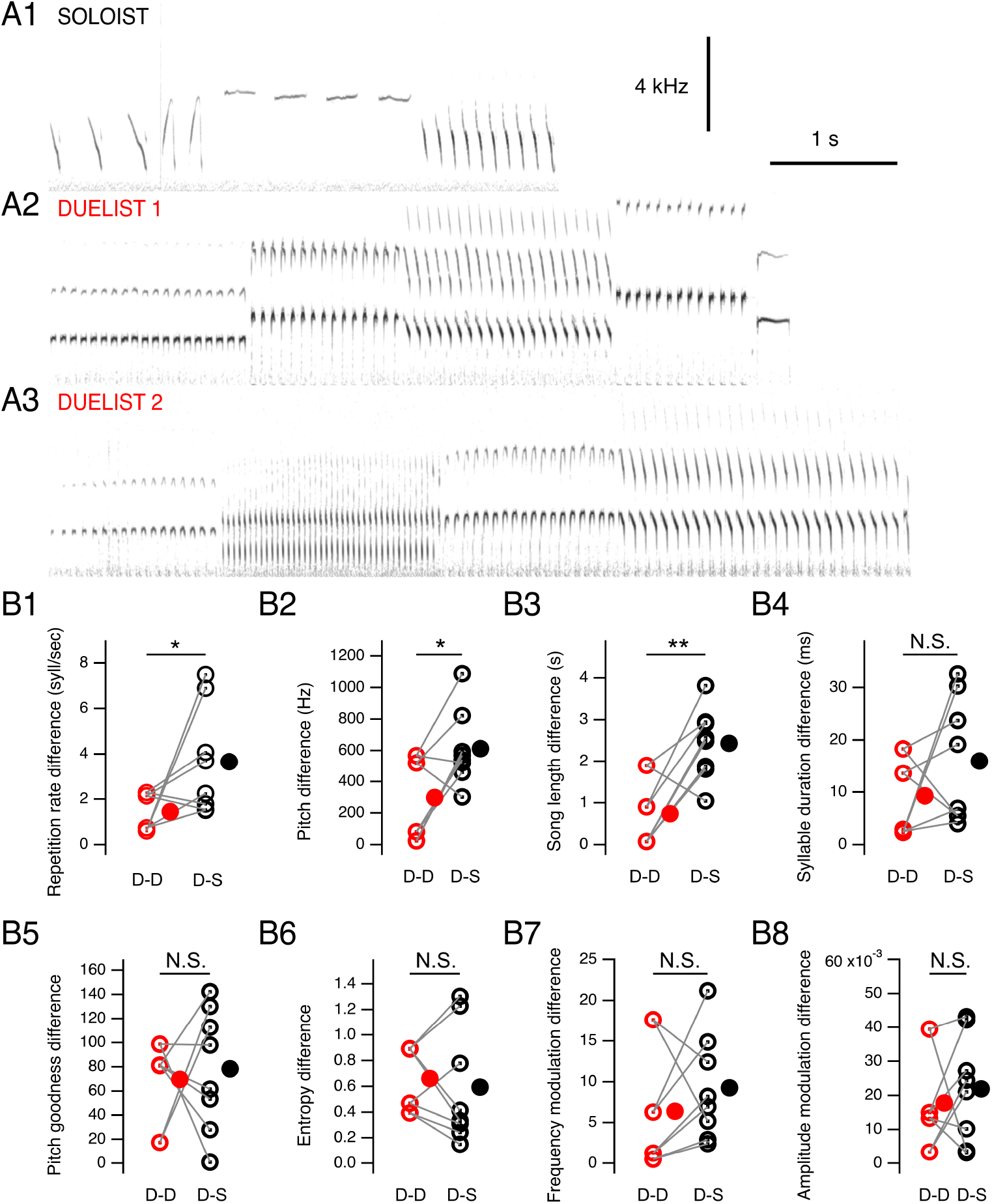
Dueling singers sing similar songs. **A.** Representative example of songs from the three canaries in a group: the ‘soloist (A1) and the two main duelists (A2, A3). **B1-B8.** Song features are compared between duelists (‘D’) and soloists (‘S’), showing more similar values between the two main duelists of each group than between each duelist and the predominantly soloist canary for song repetition rate, pitch and song length, but not for syllable duration, pitch goodness, entropy, frequency modulation and amplitude modulation of songs. Open symbols, average from each individual bird. Filled symbols, average across all birds. Differences between duelists in red and between duelists and soloists in black. * P < 0.05 and ** P < 0.01, two-tail Wilcoxon Signed-Rank test.

The average difference in syllable repetition rate was of 1.4 syllables/second between the two main duelists’ songs and 3.7 syllables/second between songs of each duelist and the soloist (Fig. 5B1, P = 0.0195, one-tail Wilcoxon Signed-Rank Test, n = 8 paired comparisons, 12 birds from 4 groups, total of 2750 songs). Similarly, the average difference in song pitch was of 298 Hz between the two main duelists songs and 612 Hz between songs of each duelist and the third bird (Fig. 5B2, P = 0.0195, one-tail Wilcoxon Signed-Rank Test, n = 8 paired comparisons, 12 birds from 4 groups). Finally, the average difference in song length was of 0.7 s between the two main duelists’ songs and 2.4 s between songs of each duelist and the third bird (Fig. 5B3, P = 0.0078, one-tail Wilcoxon Signed-Rank Test, n = 8 paired comparisons, 12 birds from 4 groups). Additional song features (i.e., syllable duration, frequency and amplitude modulation, entropy and pitch goodness) tested were not more similar between dueling birds than between dueling birds and the third bird (P = 0.191, P = 0.156, P = 0.230, P = 0.680, P = 0.473 respectively, one-tail Wilcoxon Signed-Rank Test, n = 8 paired comparisons, 12 birds from 4 groups).

Since songs for this analysis were solo songs, results show that the song acoustic properties of dueling canaries are closer to each other than they are to the canary that more rarely participates to duels, independently from their properties during singing overlapping interactions.

## Discussion

### Socially-induced plasticity of crystallized songs

One major conclusion of our study is that interactive singing dynamics influence both singing activity and song properties of male canaries. Our results show that the decision of male canaries to sing depends on the decision to sing by other males embedded in a social competition network. Remarkably, song interactions evoke song plasticity: song length and its variability differ depending on singing onset relative to the other canary (Fig. 4).

We have shown that direct interactions between male canaries through singing overlaps induce extensive plasticity of the song. During song competitions, male songbirds might take turns, i.e avoid overlap (Hultsch and Todt, 1982; Pika et al., 2018) or adjust their emissions to avoid jamming, e.g. by increasing the amplitude of the signal (Hardman et al., 2017), by using song types that differ from those of the competitor (Vehrencamp et al., 2014), or by shifting the pitch, the latter under laboratory conditions (Turner and Brainard, 2007). Jamming the rival male by simultaneous singing, is, however, difficult to estimate for third parties such as listening females. Alternatively, males might try to outcompete their rivals by prolonging their signal emission to end last. Remarkably, male canaries are able to increase song length by about 50%, which means to utter about 50% more motor gestures (syllables) during the continuing performance. Likely, only males in excellent physical condition are able to do so, in relation long songs are sang by canaries only that were exposed to elevated levels of testosterone for longer periods, a sign of fitness (Heid et al., 1985; Gil and Gahr, 2002). So far ‘crystallized’ songs were known to retain the ability to undergo small plastic changes under laboratory conditions (Tumer & Brainard, 2007; Leonardo and Konishi, 1999) but even such plasticity required substantial training of the birds, much beyond the time scale of seconds on which male canaries change their songs.

A limitation and at the same time a strength of the study is the controlled number of canaries per group studied with telemetry in an aviary. Whereas this setting has allowed us to quantify singing interactions in small groups of domesticated canaries in a naturalistic social environment, it does still not fully correspond to the social environment found in their natural habitat in Macaronesia (Voigt & Leitner, 1998). There, wild canaries habitat is more extended, and natural behavior should ideally be studied in the atlantic canary, rather than in the domesticated species. Recording with telemetry from groups of canaries in the field poses major challenges that should be addressed in the future to further characterize the singing behavior of canaries beyond this study, in their natural habitat.

### Neurobiological implications

Motor behaviors are acquired throughout plastic developmental periods during which pre-motor programs are stored in the connectivity and properties of neural networks (Sillar, Combes, Ramanathan, Molinari, & Simmers, 2008). This is the case of rhythmically-activated neural networks or central pattern generators, which retain throughout adulthood the ability to be activated in specific cellular sequences while remaining plastic (Marder, O’Leary, & Shruti, 2014), ensuring the temporal control of movements. Our results shed light into the socially-induced plasticity of ‘crystallized’ songs. The behavioral plasticity of songs found here suggests that premotor networks underlying song production undergo a reconfiguration in social contexts.

The recording of song-controlling brain nuclei during canary duels will be of great interest to understand how birdsong premotor networks are reconfigured in social environments. Songs from other canaries, and not only the bird’s own song (Lehongre & Del Negro, 2009) have been shown to evoke auditory responses in nucleus HVC, that encodes timing during singing (Hahnloser et al., 2002; Liberti et al., 2016). HVC recordings during duels should elucidate how the song of overlapping con-specifics influences ongoing activity in HVC. We find that the relative timing of singing influences song properties in a song-to-song basis, suggesting that environmental cues feed into ongoing motor commands controlling singing. Interestingly, it is known that there is a powerful inhibitory gating of HVC auditory responses during singing (McCasland & Konishi, 1981; Schmidt & Konishi, 1998). However, our results suggest that this inhibitory gating may be suppressed in competitive social contexts. It would therefore be interesting to further investigate whether inhibitory gating of auditory responses in HVC is released during singing duels. Interestingly, neurons coding for song history for up to several seconds have been described in the premotor nucleus HVC from canaries (Cohen et al., 2020). These neurons have been proposed to constitute ‘hidden neural states’ that organize songs at the time scale of seconds. When a canary faces an overlap by another canary, longer-lasting song segments occurring during the remainder of the song and responsible for plasticity in song length may be encoded by such cells.

Further, the nucleus LMAN of the anterior forebrain pathway of the song control system, may also be involved in the fast song plasticity of crystallized canary songs described here. This area is known to generate variability in the song system (Alliende et al., 2017). In relation, the firing pattern of LMAN changes depending on the social context during which singing takes place in zebra finches and which correlates with small modifications of the uttered song (Kao et al., 2008).

### Ecological, evolutionary and cultural implications

Female canaries perform more frequent copulation solicitation displays when they are presented overlapping songs relative to overlapped songs (Leboucher & Pallot, 2004). Thus, female preference for overlapping songs may have exerted an evolutionary pressure on males to outlast the song of an overlapping canary, and therefore to sing longer-lasting songs. Likewise, female preference may have selected over evolutionary time scales the neural mechanisms allowing canaries to integrate social contextual information during duels and sing longer-lasting songs.

How duels impact fitness is unclear at this stage. Indeed, although females prefer overlapping singers, they also show a preference for non-fighting canaries (Amy et al., 2008). Interestingly, in our conditions, singing duels were often accompanied by physical aggression. Future experiments in the field should reveal whether physical aggression also occurs in natural conditions between dueling birds, and the biological fitness associated to different singing strategies of leaders, followers and soloists.

Results in Fig. 5 suggest that singing similarity may predispose canaries with more similar songs to perform singing duels. However, canaries may alternatively adapt fast, with few song exposures, to the acoustic properties of another canary song. Canaries may avoid (strategy of the predominantly soloist canary) or perform both song feature matching and temporal overlaps (predominant duelist’s strategy) with canaries found in the same aviary. In these lines, previous experiments on canaries suggested that social factors may influence the adjustment of songs between individuals in the same aviary (Lehongre, Lenouvel, Draganoiu, & Del Negro, 2006). Future experiments are needed to allow disambiguating different scenarios: 1) whether canaries share song acoustic properties and a syllable repertoire that predisposes them to compete, 2) whether they instead make an adaptative choice of song properties among a library of available features in a social competition context and 3) whether dueling birds both share a preexisting repertoire and acoustic properties, and further additionally actively select more similar song features. The study further raises the question of how singing duels may have shaped the evolution of song culture, both regarding song length and similarity of songs in canary populations.

## Materials and Methods

### Ethics permits

Bird housing and all experimental procedures, including the use of audio transmitters, were approved by the government of Upper Bavaria (Ethical approval ROB-55.2-1-54-2532. Vet_02-17-211) and performed according to the directives 2010/63/EU of the European parliament and of the council of 22 September 2010 on the protection of animals used for scientific purposes.

### Animals and housing conditions

A total of 30 adult domesticated canaries (six groups formed by three males and two females each) were part of this study. Animals were randomly selected from larger aviaries and housed together in 2m^*^1m^*^1m indoor aviaries or alternatively 1.68m^*^0.78m^*^1.68m boxes. In both cases, birds were not within hearing range of other birds. Birds were kept in breeding conditions under a 13/11 Light/Dark cycle (fluorescent lamps), at 24 °C and 60–70% humidity. Food (mixed seeds, and “egg food”), fresh water and cuttlebone were provided *ad libitum*, as well as nesting material and nests.

### Audio recordings on Pico Island

Two Sennheiser MKH 70 P48 directional microphones pointed in different orientations in field where canaries had been observed. Recordings were performed with a Marrantz PMD661 recorder with an audio bit depth of 24 bit.s-1. The location of the recording areas (Fig. 1) in Arieiro and São João were 38°26’29.5’’N28°28’40.6’’W and 38°25’10.9’’N28°20’05.0’’W.

### Audio recordings at the Max Planck Institute for Ornithology

The five birds from each group used for telemetric recordings were moved from larger aviaries to smaller indoor aviaries in our institute. Custom-made wireless microphones (0.6g, including battery) (Gill et al., 2016) were used for sound recording. The wireless microphone was placed on the back and fixed with an elastic band around the upper thighs of the bird. The frequency modulated radio signals were received with the communication receivers (AOR5000, AOR, Ltd., Japan). Audio signals were fed into an eight channel audio A/D converter (Fast Track Ultra 8R, Avid Technology, Inc. U.S.A.) and recorded with custom-made software.

### Video recordings

Videos used for behavioural quantifications were recorded in aviaries with a CX405 handycamcorder (Sony) with built-in microphones.

### Song analysis

Song onset and duration were manually determined on sonograms. Analysis were performed with Matlab and Igor Pro Software. Songs recorded in the field were high-pass filtered for display using Audacity software, with a cutoff frequency of 2kHz and a roll-off of 12 dB per octave. The histograms resulting from shuffled data (Fig. 2) were generated by randomizing the onsets of all songs sung in a day, keeping the lengths and number of songs the same.

We used Sound Analysis Pro 2011 software (Version 2011.104) for segmentations and analyses of vocal sounds to generate Fig. 5. The program segments the song in an automatic process and measures the sound features of each syllable including syllable duration, mean amplitude, mean pitch, mean frequency modulation, mean entropy, mean pitch goodness, mean average frequency.

Occasionally, we recorded periods of complex interactions in which birds repeatedly overlapped each other. Since these songs could be considered as both leading and following songs, they were excluded from the analysis.

A sixth group of birds recorded was excluded from analysis due to the low number of songs that did not allow us to perform statistical analysis on singing interactions.

### Statistical analysis

Data along the manuscript are provided as mean ± S.E.M unless otherwise stated. Statistics were performed with Igor Pro Software or Matlab. Tests used are specified along the manuscript.

## Acknowledgements

PA was funded by the Munich Center for Neurosciences, the Biomentoring program of the Ludwig-Maximilians-University and the Max Planck Institute for Ornithology. SM and MG were funded by the Max Planck Institute for Ornithology. We are extremely grateful to Stefan Leitner, Lisa Trost, Roswitha Brighton and animal caretakers from our Institute for their support throughout this study. We are thankful to Hannes Sagunsky and Markus Abels for technical support, and to our neighbors in Pico island for their support while performing recordings in Pico island fields. We are very grateful to Ahmed El Hady, Wolfgang Goymann, Stefan Leitner and Frederic Theunissen for comments on previous versions of this manuscript.

